# Quantitative mRNA Imaging Throughout the Entire *Drosophila* Brain

**DOI:** 10.1101/096388

**Authors:** Xi Long, Jennifer Colonell, Allan M Wong, Robert H Singer, Timothée Lionnet

## Abstract

We describe a fluorescence in situ hybridization method that permits detection of the localization and abundance of single mRNAs (smFISH) in cleared whole-mount adult *Drosophila* brains. The approach is rapid, multiplexable and does not require molecular amplification; it allows facile mRNA expression quantification with subcellular resolution on a standard confocal microscope. Using a custom Bessel Beam-Structured Illumination microscope (BB-SIM), we further demonstrate single-mRNA detection across the entire brain sample.

---

The expression of select genes at specific times in appropriate neurons is required for brain development and function. Because of the wealth of genetic tools available, *Drosophila* has emerged as an important model system to identify the gene networks underlying neuron types and functions^1–3^. Existing approaches that quantify gene expression using sequencing methods mainly rely on isolated cell populations, in which neurons are extracted from their native context. RNA fluorescent *in situ* hybridization (FISH) is a powerful technique that allows measuring mRNA levels with subcellular resolution within preserved tissue^4-7^, but has proved difficult to apply to adult *Drosophila* brains due to problems with probe penetration and autofluorescence. Here, we develop a simple and robust whole-mount RNA FISH method for adult *Drosophila* brains to directly connect gene expression with neuronal identity and functional anatomy. Whole-Mount Imaging is advantageous compared to sectioning-based methods because it requires neither challenging sample preparation (serial sectioning, labeling, mounting, imaging) nor post-imaging analysis (stitching, registration). By optimizing the processing and clearing, we demonstrate whole brain multiplexed mRNA quantification using a standard confocal microscope. By using a custom Bessel Beam-Structured illumination microscope, we achieve single mRNA detection throughout the brain.

The overview of the method is shown in Figure 1a. In order to facilitate probe penetration, we treat the tissue with 5% acetic acid^8^, and apply a high temperature (50°C) step to enhance diffusion, before lowering the temperature to 37°C for efficient hybridization. We then quench autofluorescence using sodium borohydride and optically clear the tissue with Xylene. The method requires no molecular amplification, can be performed in 2 days, ensures optimal tissue preservation (10% formamide, a 5-fold decrease from previous techniques^9^) and is compatible with immunostaining (Figure S1).

**Figure 1.**
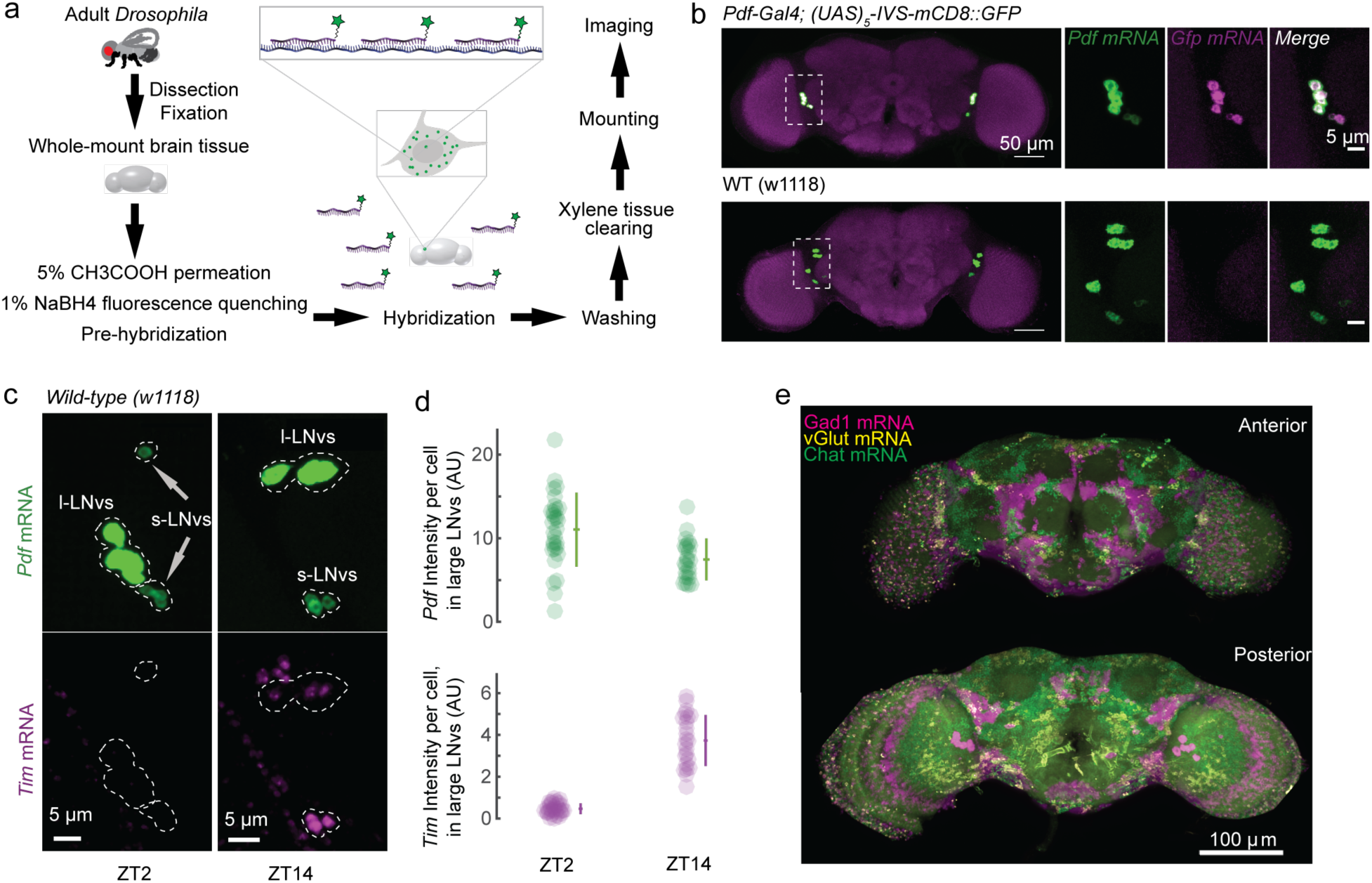
(a) Schematic illustration of the RNA FISH workflow (See Methods and Supplementary Materials for details). (b) Simultaneous detection of *Pdf* mRNA (green) and *Gfp* mRNA (magenta) using confocal microscopy. Top: maximum intensity projections of a Pdf-Gal4; (UAS)_5_-IVS-mCD8::GFP fly brain. *Pdf* and *Gfp* mRNAs are colocalized. Bottom: maximum intensity projections of a wild-type fly brain. Only the endogenous *Pdf* expression was detected; *Pdf* expression is lower in the s-LNVs (faint cells) (c) Comparison of *Tim* (magenta) expression at ZT2 (left) and ZT14 (right) in wild type, PDF neurons (green). Maximum Intensity projection of confocal microscope stacks. (d) Quantification of *Tim* expression in PDF l-LNvs neurons at ZT14 and ZT2. *Tim* expression is reduced at ZT2, while *Pdf* expression remains constant over the daily cycle. (e) Simultaneous detection of *Gad1* (magenta), *vGlut* (yellow) and *Chat* (green) mRNAs in a whole-mount *Drosophila* adult brain. Top is the anterior and bottom is the posterior view of a confocal stack (3D movie is in supplementary).

To test the specificity of our approach, we compared wild type versus transgenic flies in which Green Fluorescent Protein (GFP) is only expressed in the pigment dispersing factor (PDF) expressing neurons. PDF neurons are part of the circadian pacemaker network and consist of two groups, 4 large ventrolateral neurons (l-LNvs) and 4 small ventrolateral neurons (s-LNvs) in each hemisphere^10^. Brains labeled with FISH probes targeting GFP and PDF mRNAs in distinct colors and imaged on a confocal microscope revealed the characteristic two groups of l-LNvs and s-LNvs on each side of the brain. GFP-coding mRNA signal was not observed outside of the PDF neurons or in wild type flies, confirming specificity of the hybridization (Figure 1b).

To determine whether we could resolve changes in less abundant genes, we targeted *Timeless* (*Tim*), a transcription factor that cycles daily^11^ and regulates the circadian cycle in *Drosophila.* We acquired confocal images of light-dark cycle entrained wild type files at zeitgeber times (ZT) 2 and 14, and we observed as expected a strong *Tim* signal in PDF neurons at ZT14, while the intensity was greatly diminished at ZT2 (Figure 1c,1d and S2)^9,12^.

Because of its multiplexing potential, the technique lends itself to multiple applications. FISH can be a powerful characterization and validation tool for the rapidly expanding collections of genetic lines used in neuroscience and optogenetics. We selected three Gal4 lines (Mi1, Mi4, Mi9)^13^ with known neurotransmitter expression and crossed them with UAS-myr::HaloTag to generate expression of a HaloTag reporter in the desired neurons^14^. We profiled those brains by labeling the HaloTag protein with its fluorescent ligand in a first color (Methods) and hybridizing FISH probes targeting genes involved in distinct neurotransmitter pathways in a second color (*Gad1*, *vGlut* and *Chat*,. respectively associated with GABAergic, glutamatergic and cholinergic transmission). The overlap between the HaloTag and the FISH signals confirmed the expected neurotransmitter signatures (Figure S3). We then interrogated the simultaneous expression of three genes associated with distinct neurotransmitter pathways. Resulting confocal stacks (Figure 1e, Mov S1) display non-overlapping spatial patterns of expression, suggesting minimal co-expression of these neurotransmitters. Such multiplexing experiments provide a rapid, powerful tool to link gene expression patterns to brain-wide architecture, function and connectomics patterns.

mRNA localization has been proposed as a mechanism to regulate translation, specifically in neurons^15^. However, this model remains mostly untested in animals because single mRNA imaging in the large volumes relevant to neuronal connections *in vivo* is challenging. Hence, we sought to test the feasibility of our method to detect the location of single molecules in whole mount tissue. High-resolution imaging requires tissue transparency and matched refractive indices (RI) of the tissue and lens immersion media. To this end, we designed and built a specific Bessel Beam Selective Plane illumination microscope (SPIM) capable of structured illumination (SIM) (Figure S4). The setup was engineered to image in media with a refractive index matched to the measured index of xylene-cleared *Drosophila* tissue (Supplementary Note 1). Excitation from two directions allows SIM resolution enhancement along both X&Y (Figure 2a, Methods)^16^.

**Figure 2.**
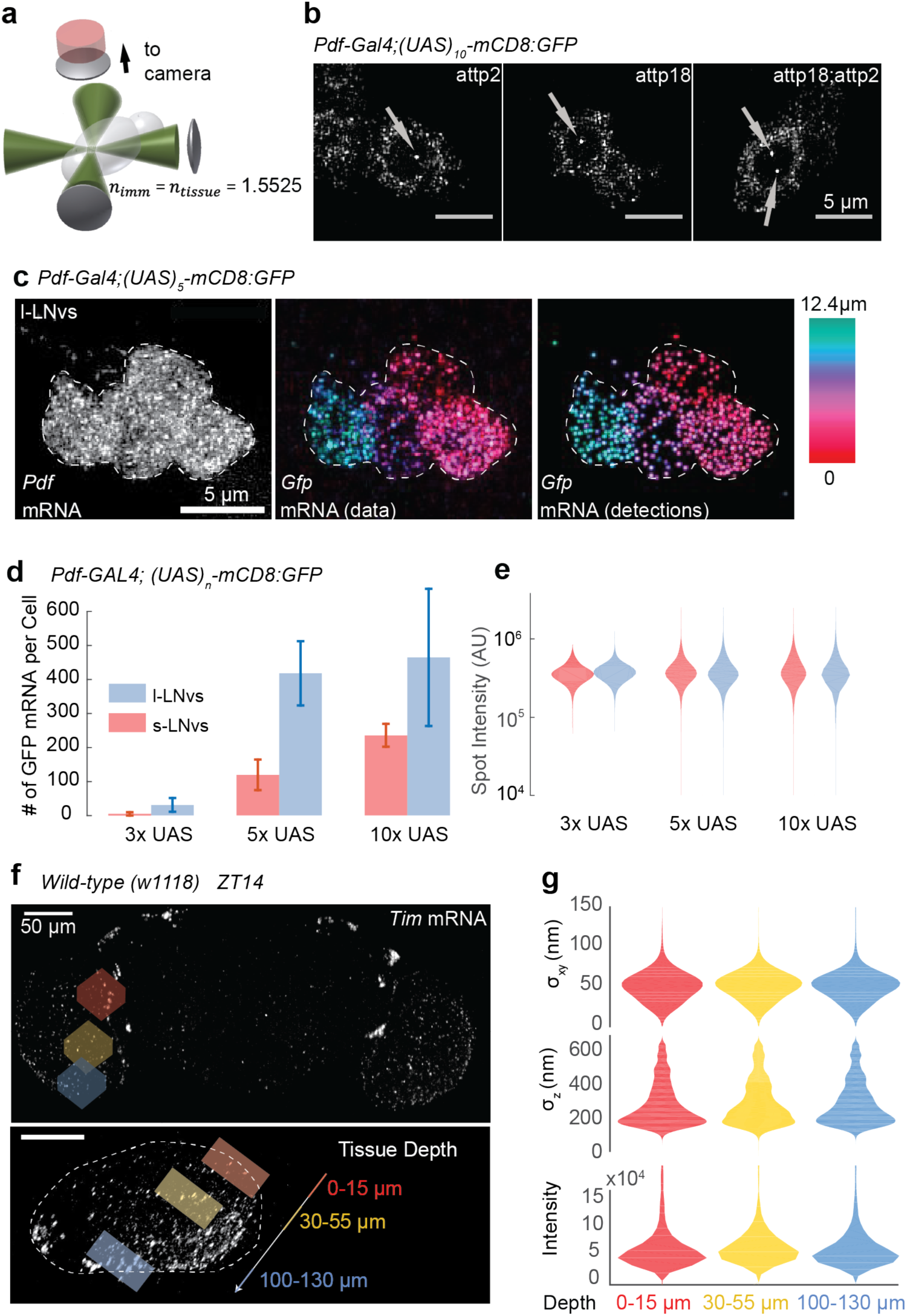
(a) Schematic of Bessel Beam Structured Illumination microscope. Imaging media has refractive index matched to xylene clearing tissue (RI=1.5525). Addition of a second excitation objective and beamline allows for Bessel Beam plane illumination for axial resolution and patterning in X&Y for lateral resolution enhancement using structured illumination. (b) PDF neurons in lines with different genomic insertions of a GFP reporter gene display a number of fluorescent nuclear foci consistent with the number of alleles. Left, Center: single genomic insertion, one nuclear focus; right: double genomic insertion, two nuclear foci. Images are Maximum Intensity Projections of confocal stacks, Transcription Sites are marked by arrows. (c) Single *Gfp* mRNAs appear as diffraction-limited spots in PDF neurons within whole-mount *Drosophila* adult brain. Left: *Pdf* mRNA signal is used as a marker for the cell bodies; center: Gfp mRNA signal color-coded for depth. Right: Reconstructed image obtained using a spot counting algorithm, color-coded for depth. (d) Number of *Gfp* mRNAs per cell in fly lines where the GFP reporter is driven by 3, 5, or 10 UAS repeats (*N =* 3,4,4 brains respectively). The colors indicate the two different PDF neuron types: blue=large PDF cells, red=small PDF cells. (e) Intensities of *Gfp* mRNA spots in 3xUAS, 5xUAS and 10xUAS lines. The colors indicate the two different PDF neuron types (*N* = 3,4,4 brains respectively) (f) Representation of the three optic lobe regions used to compare the detection performance. Images represent orthogonal maximum intensity projections of low-resolution BB-SIM scans of a brain labeled against the *Tim* mRNA. Top: anterior; bottom: sagittal view (g) Spot Intensity and Size as a function of depth in high-resolution scans of the three regions of the optic lobe labeled in panel f using probes against the *Tim* mRNA.

We first tested whether we could detect transcription sites in the whole mount brain. The multiple nascent pre-mRNAs at the locus of a transcribing gene typically generate a bright nuclear focus when imaged with FISH^4^. We imaged various lines differing in the location or number of genomic insertions of a GFP reporter gene (all under the control of a *Pdf* driver). In lines harboring a single insertion site (attp2 or attp18), we observed zero or one bright focus in each of the PDF neurons, but never observed two foci per nucleus (Figure 2b). In contrast, we frequently observed two foci per nucleus in a double insertion line (attp2; attp18). To further confirm our interpretation, we quantified the number of *Tim* and *Pdf* nuclear foci in s-LNvs, and observed a dramatic change (~5x) in the number of nuclear foci for *Tim* at ZT2 and ZT14 while the *Pdf* nuclear foci displayed little variation (Figure S5). These results agree with previous findings that *Tim* transcription is reduced at ZT2, while *Pdf* transcription does not substantially change over the daily cycle^12^.

We next set out to detect single mRNA particles. Flies with the *Pdf-Gal4* reporter expressing GFP in PDF neurons were hybridized with probes targeting the *Pdf* and the *Gfp* transcripts in separate colors, and high-resolution SPIM/SIM volumes containing the PDF neurons were acquired (Figure 2c). We detected diffraction-limited foci of the GFP-encoding mRNAs in the PDF neuron cell bodies. We could not resolve individual endogenous *Pdf* mRNAs within the cell bodies because they were too dense (left panel). Using an algorithm to characterize the individual spots^17^, the brightness of the mRNAs typically appeared as a unimodal intensity distributions above the fluorescent background (Figure S6), consistent with individual mRNAs. A characteristic of single mRNA detection is that the intensity of detected mRNAs remains constant regardless of the expression level. To demonstrate this, we used a set of driver lines with increasing number of UAS elements upstream of the GFP reporter, generating increasing levels of expression in the PDF neurons^18^(Figure S7). When imaging those brains at high resolution (Figure 2d), we counted as expected an increasing number of spots per cell with increasing number of UAS elements; importantly, the brightness of individual spots did not vary, confirming that we are detecting individual molecules (Figure 2e).

Tissue thickness can decrease detection efficiency due to aberrations and scattering. To assess the impact of tissue depth on our detection performance, we measured the distribution of fluorescence intensities and spot sizes of *Tim* mRNAs imaged in different brain regions. Spot intensities and sizes were constant through the entire brain sample, suggesting that depth had a minimal impact on detection quality and that our setup achieves diffraction-limited performance throughout the brain volume (Figure 2f,g, S6g).

We demonstrated the potential of our technique by quantifying the absolute expression of *Tim* in PDF neurons at ZT2 and ZT14 (Figure 3a,b). These results are in good agreement with confocal measurements of total intensity (Figure 1d and S2) and provide the basis for absolute gene expression studies at the scale of the entire brain.

**Figure 3.**
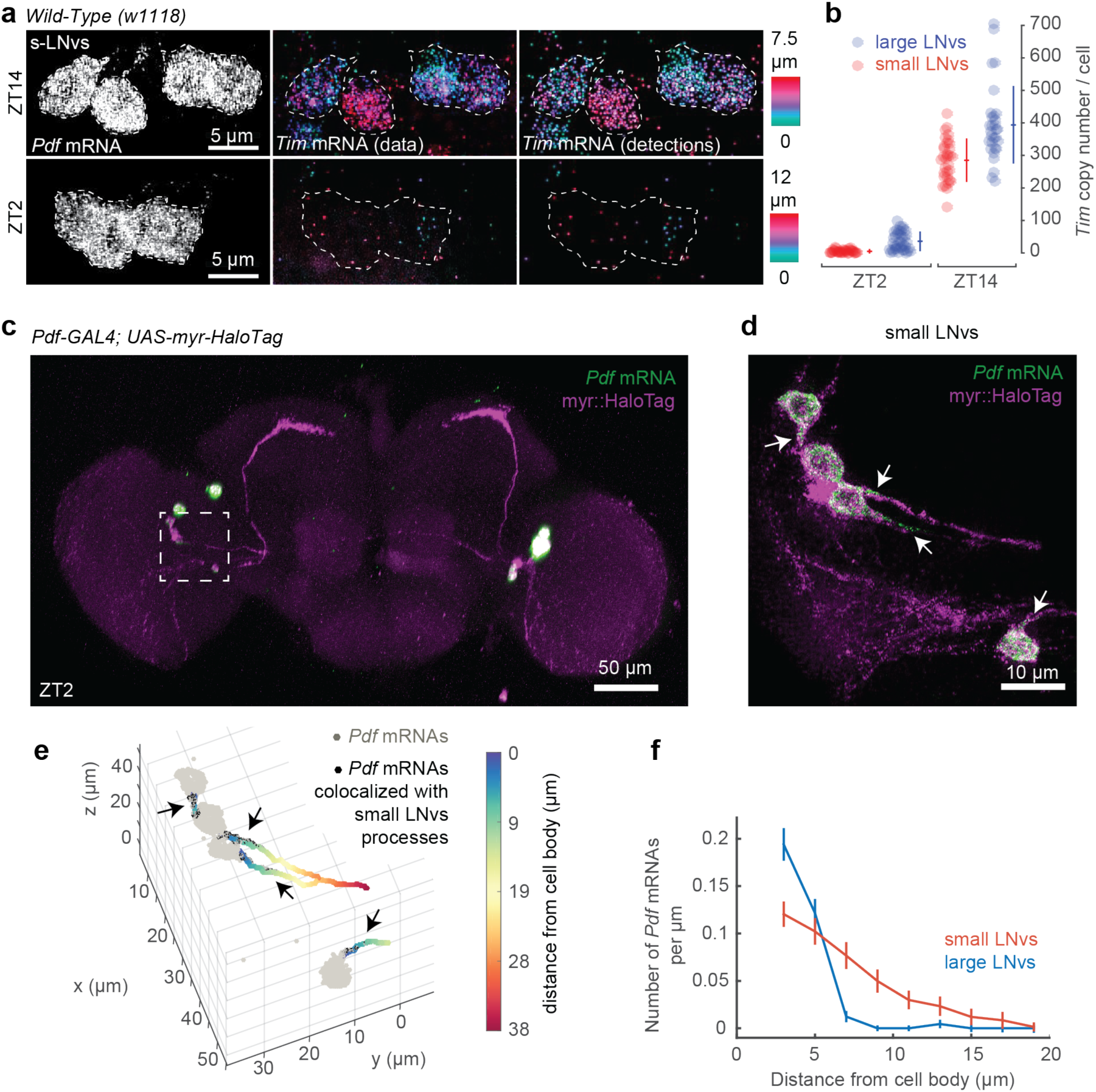
(a) PDF neurons display a large number of single *Tim* mRNAs at ZT14 (top) compared to ZT2 (bottom) in PDF s-LNvs neurons. Left: Maximum Intensity projections of BB-SIM stacks of the Pdf mRNA FISH channel are used to mark the cell bodies; Center: color-coded projections of the *Tim* mRNA FISH Channel BB-SIM stacks; Right: color-coded projection of the *Tim* mRNA image reconstruction obtained using the spot counting algorithm. (b) Quantification of the number of *Tim* mRNAs in PDF neurons at ZT14 and ZT2. The colors indicate the two different PDF neuron types. (*N*=5 brains for ZT2, 4 brains for ZT14) (c) Correlation of neuronal architecture with gene expression localization. Maximum Intensity Projection of the BB-SIM stack of a brain in which the PDF neurons are filled with a fluorescent HaloTag ligand (myr::HaloTag, purple) and the *Pdf* mRNA is imaged by FISH (green). (d) Highresolution scan of the boxed region in panel c displaying 4 s-LNvs. *Pdf* mRNAs (green) accumulate within the cell bodies and in the proximal region of the processes (arrows). (e) Representation of the localization analysis results: *Pdf* mRNAs detected are represented by light gray spots; *Pdf* mRNAs that localize with processes are indicated as black spots and their prominent locations highlighted with arrows. Processes emanating from the s-LNVs are color coded for the distance from the cell body. (f) Quantification of the density of *Pdf* mRNA molecules in PDF neuron processes as a function of distance from the cell body, showing a PDF cell type specific localization profile.

While most mRNAs localize to neuronal cell bodies, we also detected molecules in processes (Figure 3c-f). The mRNA levels decreased with increasing distance from the cell bodies (similar to mammalian neurons in culture^15^) in a cell type specific manner suggesting distinct architectures or transport regulation (Figure 3f).

The whole-mount smFISH technology enables the interrogation of gene expression levels at the single-cell level using a standard confocal microscope. Because of its simplicity and versatility (multiplexable, compatible with HaloTag labeling and immunofluorescence), we anticipate it will constitute an important tool for addressing the cellular and molecular basis of brain function. Furthermore, when combined with high-resolution microscopy, it is able of detecting and counting individual molecules within an intact entire brain. This permits visualizing neuronal architecture and connections simultaneously with the levels and localization of mRNA expression. Combined with the powerful genetic tools inherent to *Drosophila*,. this technique constitutes a unique tool to address questions such as the role of mRNA localization in memory formation. Expansion of the technique, e.g. multiplexing many more mRNAs^7,19^, will further exploit its potential.

## Acknowledgements

We thank members of the former Transcription Imaging Consortium, M. Rosbash, L. Lavis, T. Brown, U. Heberlein and Y. Wu for valuable suggestions. We are grateful to A. Nern for providing the Mi4, Mi1 and Mi9 transgenic flies, G. Henry and F. Davis for providing transcriptome profiling information, and G. Ihrke and K. Close for helping *Drosophila* dissection. We also thank E. Betzig for consultation on the design of the BB-SIM microscope and the control software for the system; L. Shao for structured illumination analysis code; A. Taylor for assistance with confocal imaging. Funding for this work was provided by the Howard Hughes Medical Institute.

## Author contributions

XL and TL designed the experiments and wrote the manuscript. XL performed the experiments and did confocal measurements. JC designed and built the Bessel Beam SPIM with SIM microscope, and performed measurements using this microscope. XL, JC and TL analyzed the data. AW created the transgenic flies for detecting 2 GFP transcription sites and contributed to the neurotransmitter experiments. RS consulted on the research and helped to write the manuscript. All authors edited the manuscript.

## ONLINE METHODS

### *Drosophila* brain tissue and preparation

Pdf-GAL4 was from J. Park; pJFRC4-3XUAS-IVS-mCD8::GFP, pJFRC5-5XUAS-IVS-mCD8::GFP and pJFRC2-10XUAS-IVS-mCD8::GFP were from Pfeiffer 2010. Mi1 (*55C05*-*p65ADZp*(*attP40*); *71D01*-*ZpGdbd*(*attP2*)), Mi4 (*48A07*-*p65ADZp*(*attP40*); *79H02*-*ZpGdbd*(*attP2*)) and Mi9 (*48A07*-*p65ADZp*(*attP40*); *VT046779-ZpGdbd*(*attP2*)) were provided by A. Nern. UAS-myr-HaloTag was from Kohl 2014. We used w1118 wild-type for timeless experiments. To generate the PDF- GFP titration flies, we crossed Pdf-GAL4 to stocks that contained 3x, 5x, or 10x UAS-IVS-mCD8::GFP. For the double GFP insertion line, we made a stock of pJFRC2-10XUAS-IVS-mCD8::GFP(attp18);;pJFRC2-10XUAS-IVS-mCD8::GFP(attp2). The stock was then crossed to Pdf-Gal4. To generate selective expression of a HaloTag reporter in specific neurons of the optic lobe, we crossed Mi1, Mi4, Mi9 to UAS-myr::HaloTag. Flies were reared on standard cornmeal/agar media supplemented with yeast under 12 h light/12 h dark cycles at 25 °C. 3-5 days old adult flies were collected, then dissected in phosphate-buffered saline (PBS) and fixed in 2% paraformaldehyde for 55 min at 25 °C under normal lighting. For ZT14 samples, brain tissues were dissected under red LED lighting. The brain tissues underwent dehydration before they were stored in 100% EtOH overnight.

### FISH

Amino-labeled oligonucleotide probes^5^ (Biosearch Technologies) were labeled to NHS-ester fluorophores. To perform FISH, rehydrated *Drosophila* brain tissues were exposed to 5% acetic acid at 4°C for 5 min. After being fixed in 2% paraformaldehyde for 55 min at 25 °C, the tissues were incubated in 1xPBS with 1% of NaBH_4_ at 4°C for 30 min, followed by a 2 hr incubation in pre-hybridization buffer (15% formamide, 2x SSC, 0.1% triton X-100) at 50 °C. The brain tissues were transferred to 50uL of hybridization buffer (10% Formamide, 2xSSC, 5xDenhard’s solution, 1mg/ml Yeast tRNA, 100ug/ml, salmon sperm DNA, 0.1% SDS) with FISH probes (50-100ng/uL per reaction, containing probe sets against one or multiple genes) and incubated at 50 °C for 10 hr, followed by an additional 10 hr incubation at 37 °C. After a series of wash steps, the brain tissues were dehydrated and introduced to xylene for tissue clearing. A detailed description of the FISH protocol is included in the supplementary information.

### HaloTag staining and IHC

The brains of flies expressing the HaloTag reporter were stained with HaloTag-JF646 during fixation. After dissection, brain tissues were transferred to 2% paraformaldehyde containing 2uM of HaloTag-JF646^20^. They were then incubated with agitation for 55 min at 25 °C, followed by a series of 1xPBS washes. Antibody staining was performed after the FISH labeling. After a series of wash steps, the brain tissues were blocked with 10% normal goat serum in PBT at room temperature for 2 hr, followed by overnight primary antibody staining (Rabbit polyclonal anti-GFP Fraction, Life Technologies A11122, 1:1000 dilution with 5% NGS) at 4°C. After washing the primary antibody, brain tissues were incubated in secondary antibody (AF488 Goat anti-rabbit, Invitrogen A11034, 1:1000 dilution with 5% NGS) overnight at 4°C. Brain tissues were then fixed in 2% paraformaldehyde for 55 min at room temperature before dehydration and xylene clearing.

### Confocal Imaging

For confocal imaging, all brain tissues were attached to poly-L-lysine coated cover glass and mounted in DPX (Janelia Adult *Drosophila* CNS DPX mounting protocol). We used a Zeiss LSM 880 confocal microscope (561 nm and 633 nm laser lines) imaged through a 25X DIC NA 0.8 oil objective to detect GFP transcripts in Pdf-Gal4;JFRC5 constructs and wild type flies (Figure 1). Images were acquired with sequential excitation as stacks with 1.5 um z-spacing. Detector gain and laser power were kept constant for all samples. To detect *Gfp* transcripts in the UAS titration experiments, we used a Zeiss LSM 800 confocal microscope (561 nm and 633 nm laser lines) imaged through a 63 X DIC NA 1.4 oil objective. Images were acquired with sequential excitation as stacks with 1.0um z-spacing. Detector gain and laser power were kept constant for all samples. To detect the overlapping neurotransmitter expression in Gal4 lines, we used a Zeiss LSM 880 confocal microscope (561 nm and 633 nm laser lines) imaged through a 63X DIC NA 1.4 oil objective. Images were acquired with sequential excitation as stacks with 1.00 um z-spacing. The multiplex neurotransmitter images were acquired on a Zeiss LSM 880 confocal microscope (488 nm, 561nm and 633 nm laser lines) imaged through a 25 X DIC NA 0.8 oil objective. Images were acquired with hyperspectral lambda detection scan as stacks with 1.00 um z-spacing. Summary of imaging parameters are in Supplementary Table 1.

### SPIM/SIM Imaging

To generate the SPIM/SIM images, the brain tissue was mounted on a 1.5x3mm poly-L-lysine coated coverslip attached to the end of a 30mm glass rod. The objectives and sample were all immersed in the imaging medium. The images in this work were acquired with a NA_Bessel max_ of 0.62, which creates a Bessel beam with a central peak width 0.35μm for 560 nm illumination. 11 phases were collected for each excitation direction. The exposure time for each image was 20 msec. The lateral field of view, which is determined by the length of the Bessel beam, was 35x35μm. Z stacks up to 24μm were acquired by moving the sample with a piezo stage with closed loop position control. Larger volumes were covered by tiling, moving the sample with piezo inertia drives in XY&Z. For more details see Supplementary Note 1, Figure S4 and Supplementary Table 1.

### Image Analysis

Image Analysis Details are discussed in Supplementary Note 3.

**Figure S1.**
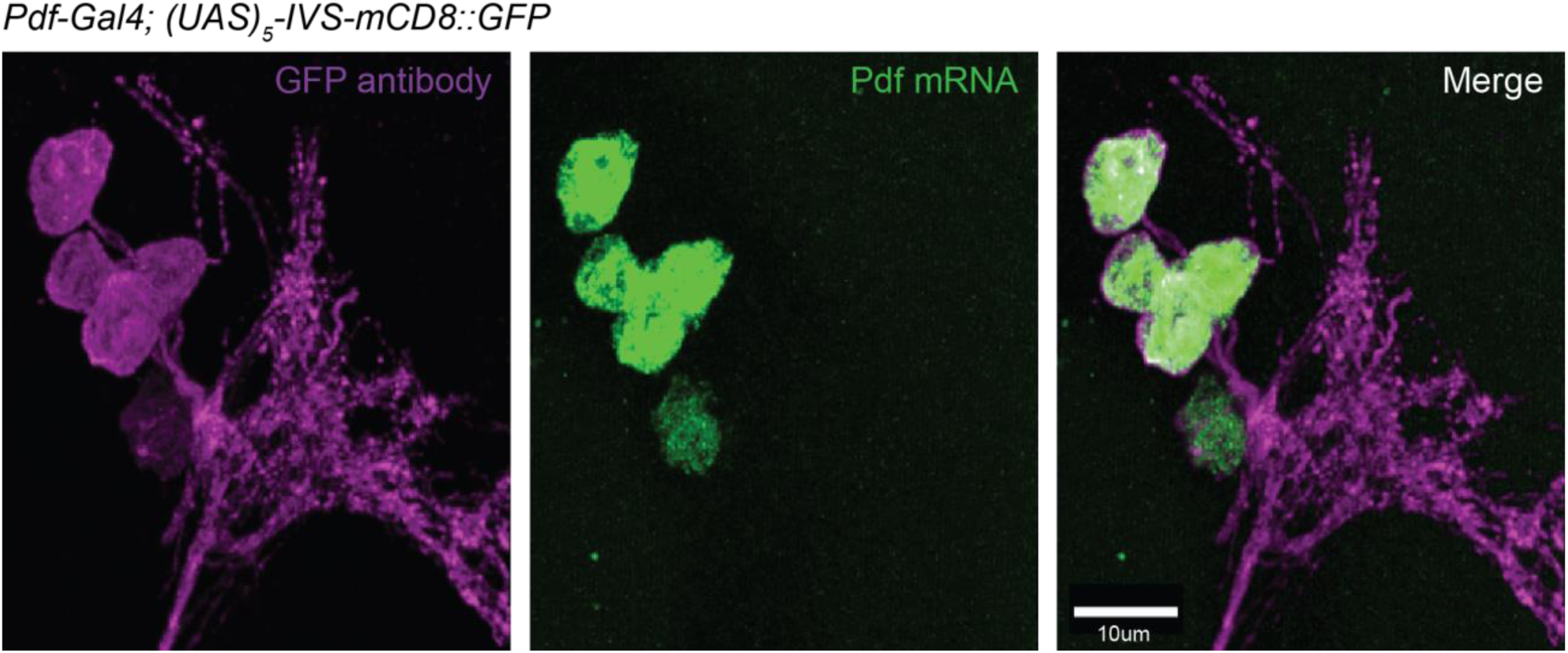
Simultaneous imaging of *Pdf* mRNA (green) and GFP protein (purple) in whole-mount tissue. Maximum intensity projection of confocal stack of a Pdf-Gal4; 5xUAS-IVS-mCD8::GFP fly brain. *Pdf* mRNA (labeled with FISH probes) accumulates mainly in the cell bodies of PDF neurons identified by the GFP protein signal (immunohistochemistry with antibodies against GFP, see Methods).

**Figure S2.**
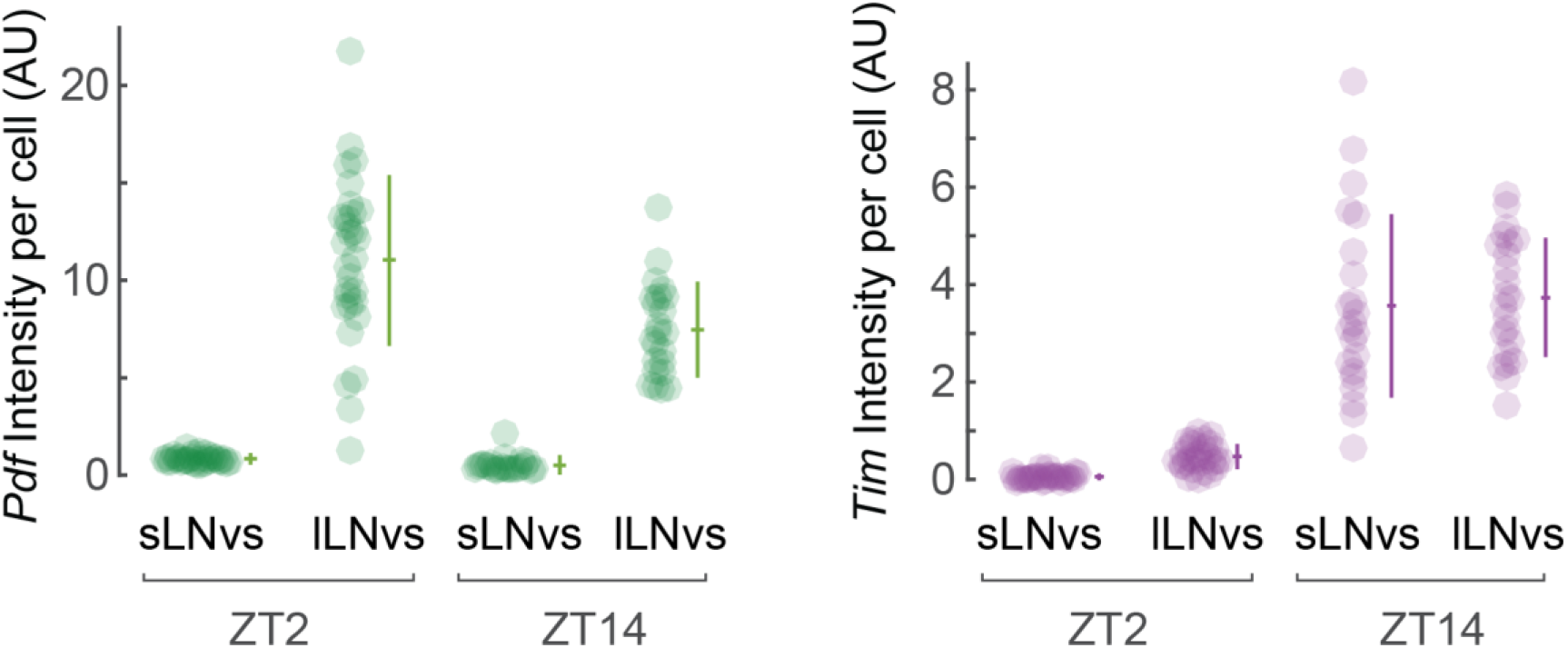
Quantitative analysis of *Tim* expression in PDF neurons (l-LNvs and s-LNvs) at ZT2 and ZT14. Cell Intensities were calculated from confocal stacks of wild type flies labeled with FISH probes against *Tim* and *Pdf* mRNAs. *Tim* expression is reduced at ZT2, while *Pdf* expression displays only modest changes over the daily cycle. Each circle corresponds to one cell; Mean and Standard Deviation are represented next to each bee swarm plot; 3 brains were imaged for ZT2, 4 brains for ZT14.

**Figure S3.**
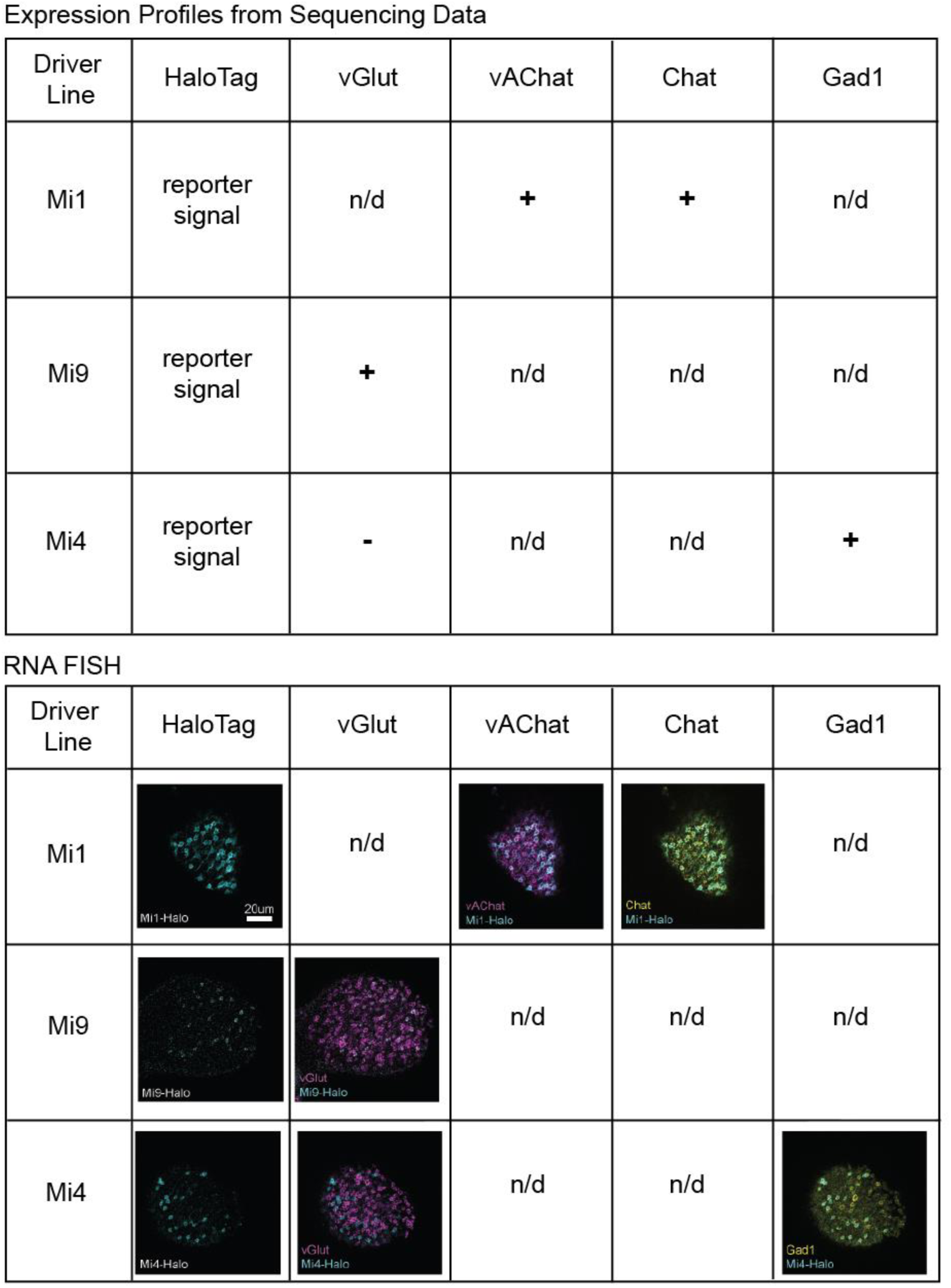
RNA FISH expression patterns confirm results previously validated by sequencing. Top: Summary of RNA sequencing data obtained in purified cells from the following optic lobe neuron types: Mi1, Mi4, Mi9 (Strother et al., Submitted). + indicates gene expressed, -indicates no expression, n/d-genes not tested by FISH. Bottom: representative confocal sections of the optic lobe from each driver line; all reporter lines express a HaloTag reporter under the control of Gal4. HaloTag fluorescent ligand (cyan); FISH probes targeting the following genes: *vGlut* (magenta), *vAChat* (magenta), *Chat* (yellow) and *Gad1* (yellow). The RNA FISH overlapping with the HaloTag signal follows the patterns predicted by the RNA sequencing data.

**Figure S4.**
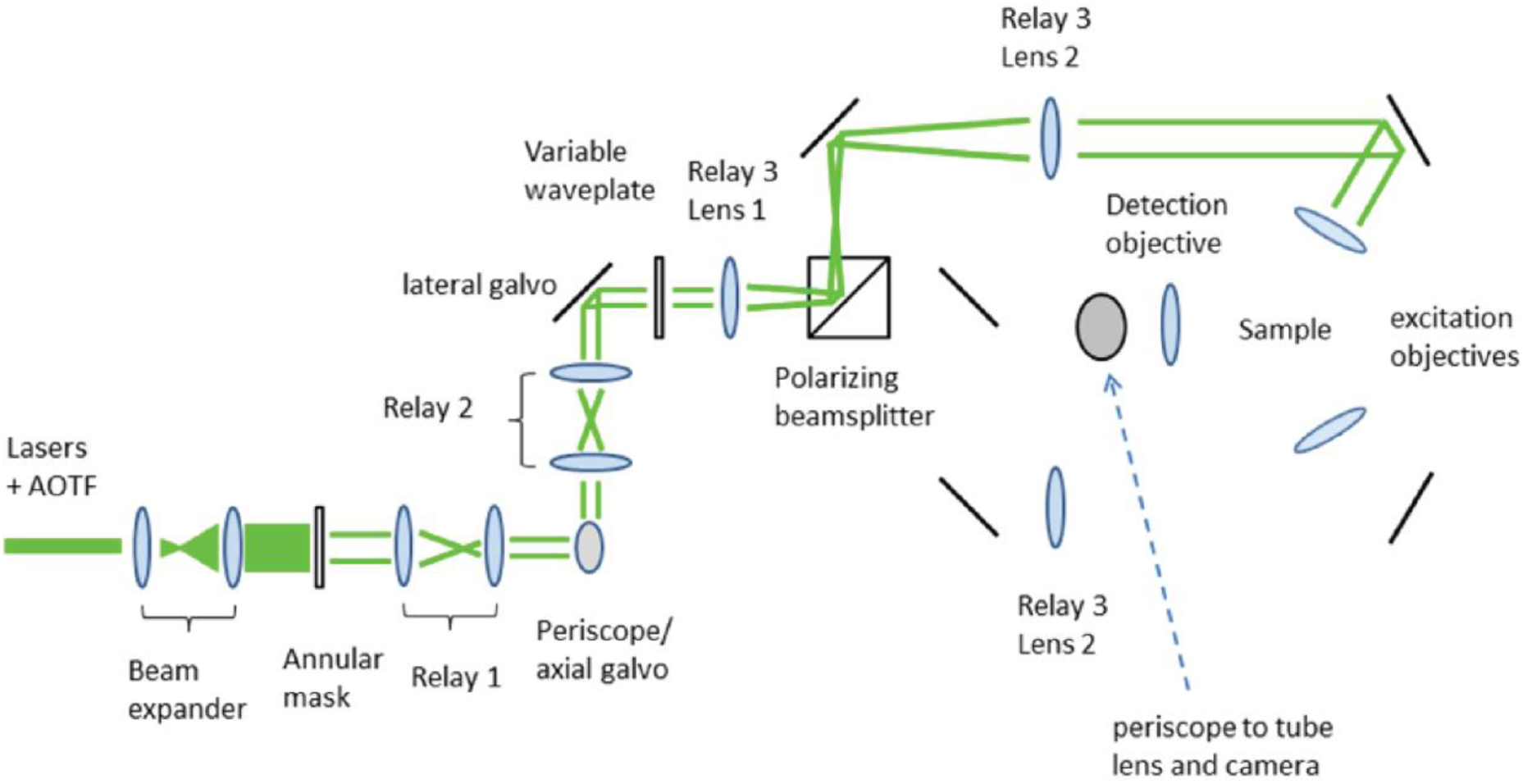
Simplified optical layout of the SPIM/SIM microscope. Beams from three lasers (488 nm, 561 nm, 640 nm) are expanded to a diameter of 3 mm, and vertical polarization set with a half wave plate. This combined beam is sent through an Acousto-optic Tunable Filter (AOTF, AA Optoelectronic) and beam expander. From the beam expander, the beam passes through an annular apodization mask (made in house using a laser mill to ablate the desired pattern on a Thorlabs Neutral Density filter) and a pair of galvo mirrors (Cambridge Technology) that allow lateral and axial positioning of the beam. The combination of a liquid crystal variable waveplate (Edmund Optics) and polarizing beamsplitter allows the beam to be directed to either excitation objective; this diagram shows the beam for just one path. The annular mask, X galvo, Z galvo, and back aperture of the excitation objectives are all at conjugate planes, so beam position is constant at those points. When imaging, the illumination plane and detection objective are fixed, and the sample is moved to image through Z to create a volume.

**Figure S5.**
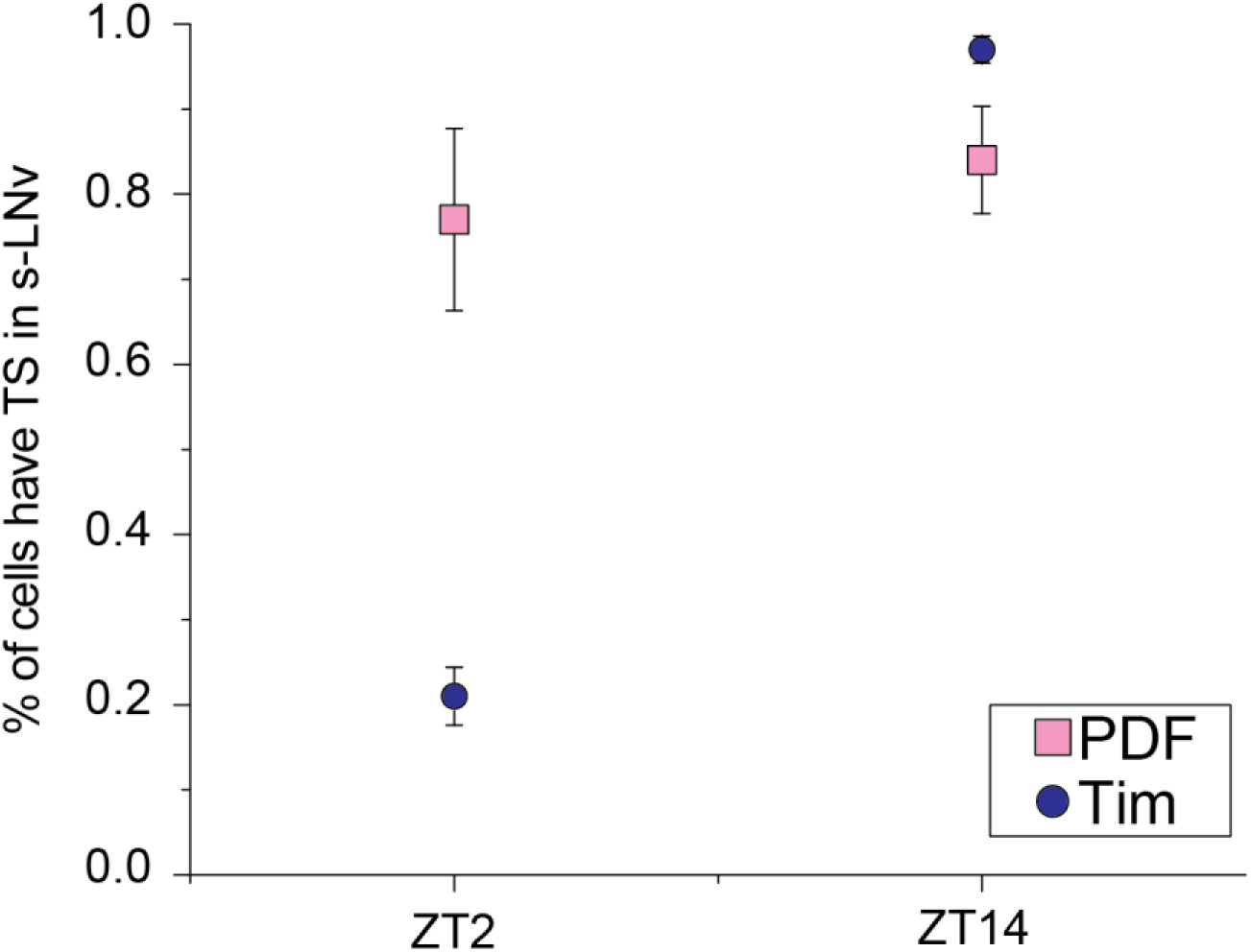
Comparison of the number of *Pdf* (pink) and *Tim* (blue) transcription sites at ZT2 and ZT14 in s-LNvs in wild type flies imaged with the BB-SIM microscope. The variation of *Tim* and *Pdf* nuclear foci is in good agreement with previous findings that *Tim* transcription is reduced at ZT2, while *Pdf* transcription remains constant over the daily cycle. 5 brains were imaged for *Tim* ZT2, 4 brains for *Tim* ZT14; 4 brains were imaged for *Pdf* ZT2 and ZT14;

**Figure S6.**
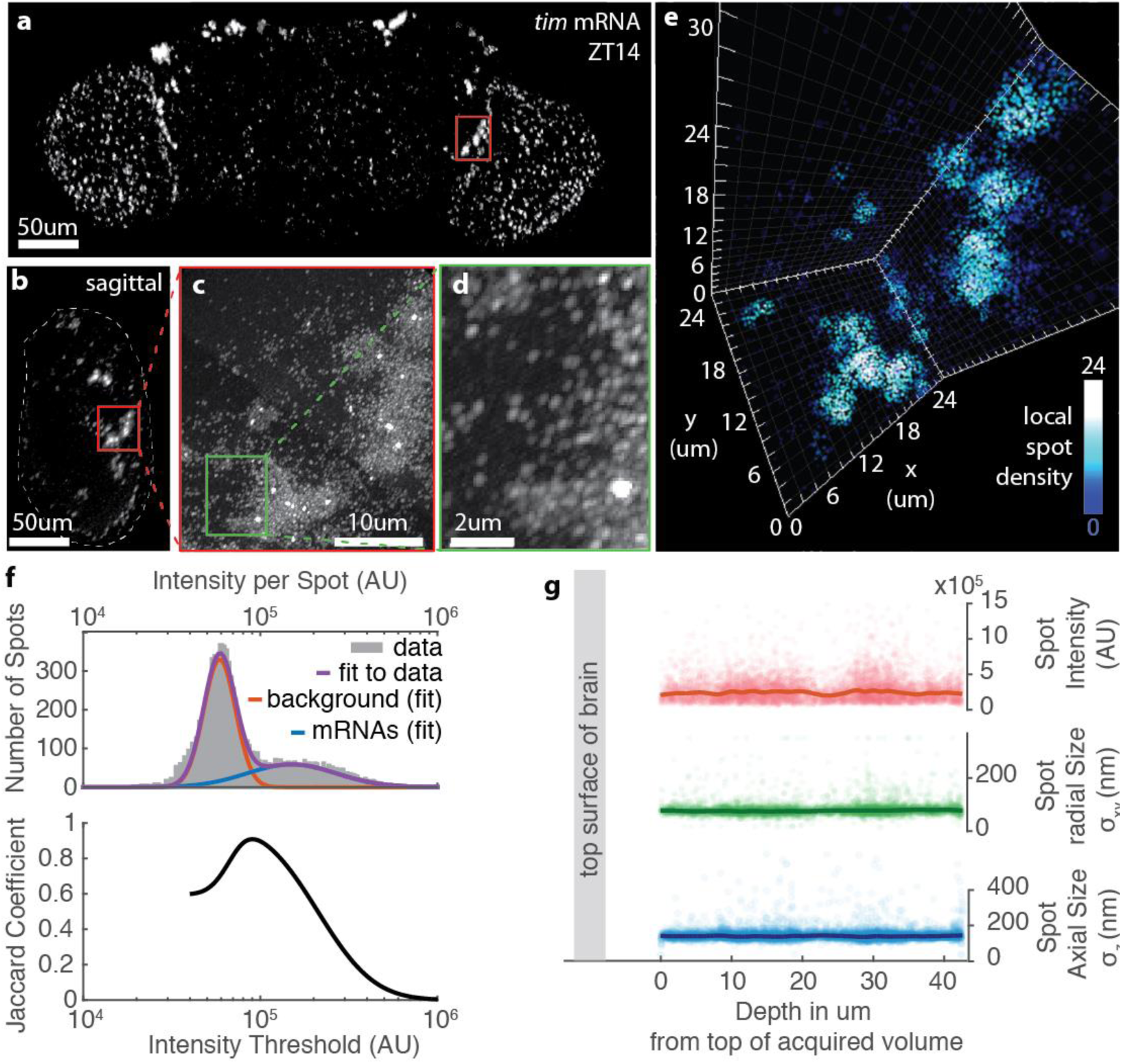
Single molecule detection of *Tim* mRNA. **a)** Low-resolution frontal view of *Tim* mRNA detected with the BB-SIM microscope in a whole-mount wild type fly (ZT14). Red box indicates the location of the right hemisphere PDF neurons. **b)** Sagittal view of the sample in a. **c)** Maximum Intensity Projection of a high resolution BB-SIM scan of the region of the PDF neurons (red box in b). High levels of expression are visible in the 8 PDF neurons; lower *Tim* expression is also present in surrounding cells. Bright foci are observed at the sites of active transcription in the nuclei. **d)** magnification of panel c displaying individual mRNA spots as well as one transcription site. Individual mRNAs appear as oblong oblique spots because the light sheet observation angle is inclined at a ~45° angle relative to the vertical axis of the image. **e)** 3D visualization of the region surrounding the PDF neurons (red box in b panel or entire c panel). The color encodes the local density of spots. **f)** Top: Histogram of the intensity distribution for the spots detected in panel c using a low detection threshold (gray bars). Background detections appear as a low intensity spots while mRNAs accumulate in a distinct high-intensity distribution. The fit to a 2-component lognormal distribution (purple) provides the respective contributions of background (red) and mRNAs (blue) to the overall histogram. Bottom: Jaccard Coefficient quantifying the similarity between the detected spots and the expected mRNA counts, as a function of an arbitrary intensity threshold (the Jaccard calculation is based on the 2-component lognormal fit, see Supplementary Note). The Jaccard Coefficient provides a metric of detection accuracy taking into account both sensitivity and selectivity; it reaches a maximum for the intensity threshold that separates optimally the background from mRNA signal. The maximum value indicates that the spots detected at the optimum are ~90% similar to the original mRNAs. **g)** Intensity and sizes of *Tim* mRNA spots as a function of depth. The fluorescence intensities and spot sizes of *Tim* mRNAs detected in panel c do not change as a function of depth. Each circle represents one detected spot; the line is a sliding average.

**Figure S7.**
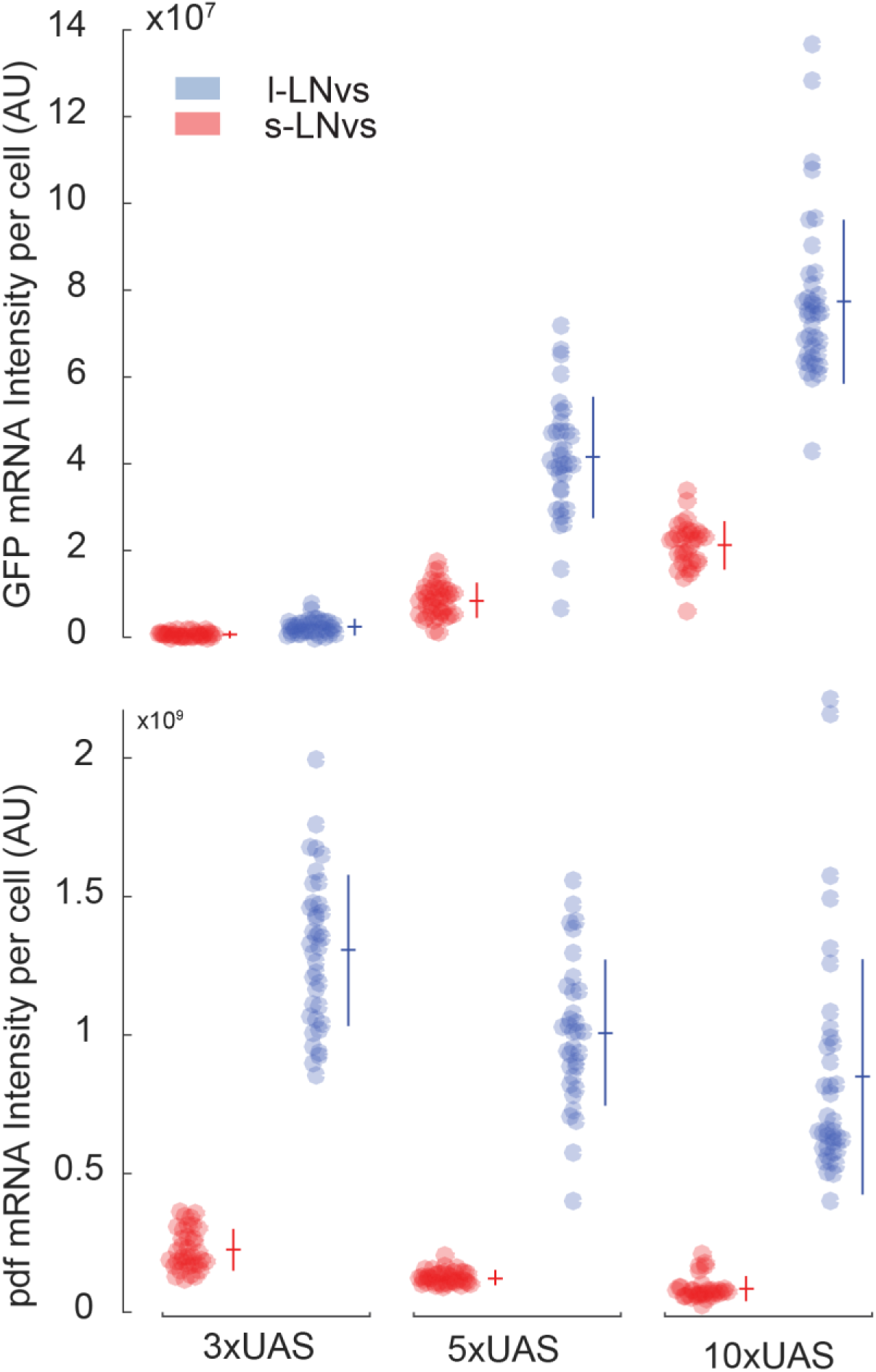
Top: comparison of *Gfp* mRNA fluorescence intensity per cell in fly lines using confocal microscopy. The GFP reporter is driven by 3, 5, or 10 UAS repeats, resulting in a gradual increase in expression of the *Gfp* mRNA. In contrast, the endogenous *Pdf* mRNA levels do not increase with increasing UAS number (bottom). The colors indicate the two different PDF neuron types. Each circle corresponds to one cell; Mean and Standard Deviation are represented next to each bee swarm plot; 5 brains were imaged for each condition.

**Supplementary Table 1.**
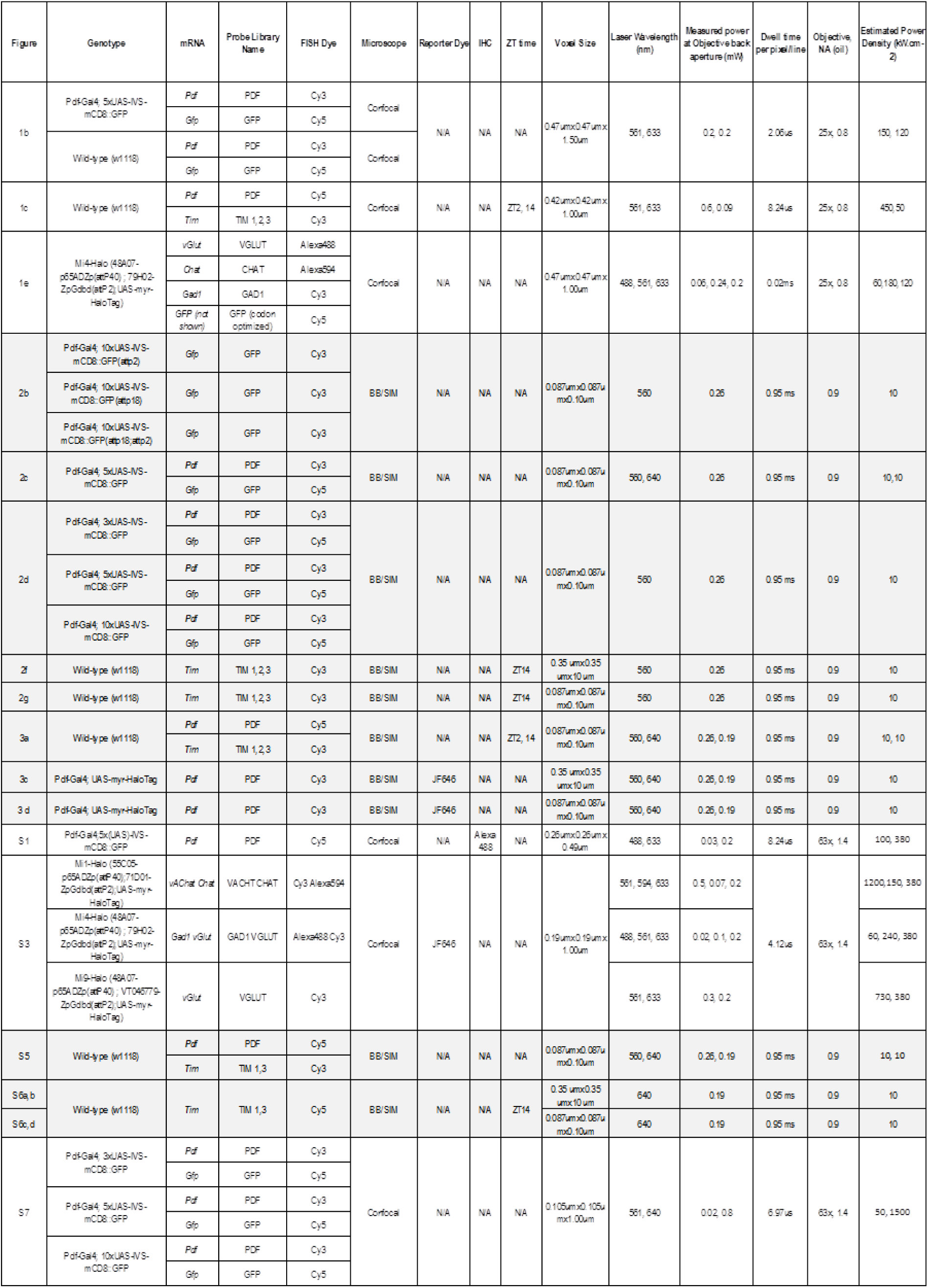
(Summary of experimental parameters for each Figure)

## Supplementary Note 1: BB/SIM microscope

The SPIM/SIM microscope is closely related to the Bessel Beam Plane illumination microscope described by Planchon, et al.^1^ and Gao, et al.^4^, which uses a Bessel beam, with a very narrow central peak (0.3 μm) and long axial extent (30-50 μm), swept or stepped to illuminate a thin sheet in the sample. As is generally true for plane illumination microscopy, the lateral resolution is limited by the lateral resolution of the detection objective, while the axial resolution is largely determined by the thickness of the light sheet. Stepping the beam creates a virtual grating: stripes perpendicular to the scan direction. A set of images collected with different positions of the grating can be analyzed using Structured Illumination (SI) analysis^3,4^ to extract high frequency information and reconstruct the image with lateral resolution twice that of the detection objective. However, that resolution enhancement is limited to the direction perpendicular to the grating. Imaging whole mount Drosophila brains with nearly isotropic resolution requires two new developments: creation of optically clear samples and illumination patterning in two directions for SI resolution enhancement in both X&Y. As shown in Figure 1, the microscope therefore has three objectives: the imaging objective plus two orthogonally mounted excitation objectives. For aberration free imaging, the system must be designed for media with refractive index precisely matched to the cleared tissue.

The first step is development of an immersion fluid precisely matched to the refractive index of xylene cleared Drosophila tissue. Dehydration of tissue followed by treatment with xylene or other high index solvents to create optically clear samples is an old technique (see, for example^2^). Such samples are usually mounted in a solid medium between a slide and coverslip. These samples are durable, but the geometry does not allow sufficient optical access to allow plane illumination, and the refractive index of the material surrounding the tissue is poorly controlled. A liquid imaging medium allows tuning of the refractive index to match the tissue and a dipping imaging geometry compatible with selective plane illumination.

The refractive index of the tissue was determined by imaging known targets - fluorescent beads—through cleared tissue in test media using a standard confocal microscope. Refractive index mismatch between the tissue index and the surrounding medium creates lensing effects which can be measured in the bead images. The composition of the test medium was varied systematically to minimize effects on the imaging. Best results were achieved with the refractive index of the medium at 1.5525. The other requirements for the imaging medium are: miscibility with xylene, low fluorescence, and relatively low vapor pressure. To meet these, the system is designed for imaging in a mixture of 90% 1,2-dichlorobenzene and 10% 1,2,4-trichlorobenzene.

The detection objective and two excitation objectives are custom designed (Special Optics/Navitar) with: working distance = 3.0 mm; design NA = 1.0 (measured NA 0.9-0.93); optical match to the index of the medium; small physical package allow 3 objectives to be mounted orthogonally with a common focal point; and solvent resistance. To allow adequate optical access to the sample, the tissue is mounted on a 1.5x3mm coverslip attached to the end of a 30 mm glass rod which holds the sample at the focus of the three objectives. The objectives and sample are all immersed in the imaging medium. A simplified optical layout of the system is given in supplementary figure M1.

The control electronics and software are identical to those described in Planchon, et al^1^. and Gao, et al.^4^, with the addition of control of the liquid crystal variable retarder to switch between beamlines. Images are collected by stepping the excitation beam along one axis with the camera exposing through all steps. The step size is (((*N-1*)/*2*)\**λ*/(*2*\**NA_Bessel max_*)) where *N* is the number of phases to be collected. That image is saved, and the imaging is repeated for all phases, with the pattern phase shifted by *2π*/*N* relative to the last. This process is repeated using excitation through the second beamline, which creates an image with patterning in the orthogonal direction. The sample is then moved in z, and the next plane imaged in the same way. For two color images, the full set of phases is collected for first color, then a full set of phases for the second color, before moving the sample and imaging the next plane. The two volumes created by imaging with X and Y modulation can show 100-200 nm displacement in z from each other, due imperfect alignment of the beams and beam deflection due to residual index variation in the tissue. Therefore, the “X image” is aligned in Z to the “Y image” before combining the two directions in the structured illumination analysis. The structured illumination analysis algorithm is detailed in Gustafsson, et al.^3^; modifications specific to Bessel beam imaging are discussed in Gao, et al^4^. The reconstructed images in different colors are registered to each other by applying x, y and z offsets measured on beads labeled with both colors.

The images in this work were acquired with an excitation aperture *NA_Bessel max_* of 0.62, which creates a Bessel beam with a central peak width 0.35 μm with 560 nm illumination. 11 phases were collected for each excitation direction. The exposure time for each phase is 20 msec; collection of one z slice in one color (22 images) takes 1.5 sec. PSF measurements with these conditions on 150 nm beads yield a lateral fwhm of 0.20 μm, and axial fwhm of 0.29 μm. The XY field of view, which is limited by the axial extent of the Bessel beam, was 35x35μm. Z stacks up to 24 μm are acquired by moving the sample with a piezo stage with closed loop position control (Physik Instrumente P-753). Larger volumes are covered by tiling, moving the sample with piezo inertia drives (Physik Instrumente, LPS-45) in XY&Z. Light collected by the detection objective is imaged through a 400 mm achromat (Edmund) onto the camera (Hammamatsu Orca 4.0).

## Supplementary Note 2: Stepwise Protocol for whole-mount RNA FISH of *Drosophila* adult brain

### Probe Library Design and Labeling

FISH probe libraries were designed based on transcript sequences using the online Stellaris Designer and purchased from Biosearch Technologies. Libraries for each gene typically consist of ~50 probes, but we were able to detect mRNAs with as little as 20 probes (e.g. in the case of *Pdf)* when the mRNA length or sequence composition did not permit designing 50 probes; increasing the number of probes increases the signal to noise ratio. Each probe is 18-22nt long with a 3’ end amine-modified nucleotide that we directly couple to an NHS-ester dye according to the manufacturer’s instructions (Life Technologies). Excess dyes were added to the reaction to ensure all probes were coupled to dyes. To separate the free dyes from dye-coupled oligos, we used the Qiagen Nucleotide Removal Columns. This approach yields 85-100% of dye-labeling efficiency. We validated our approach using HPLC purification. Probe library sequences are listed in Supplementary Table 2.

### FISH Staining

#### Solution

1xPBS

5%PBT (1xPBS, 0.5% Triton)

5% (v/v) CH_3_COOH

1xPBS with 1% NaBH_4_

Pre-hybr soln (15% formamide, 2x SSC, 0.1% triton)

Hybr soln (10% Formamide, 2x SSC, 5x Denhard’s soln,1mg/ml Yeast tRNA,100ug/ml, Salmon sperm DNA, 0.1% SDS)

30% formamide washing soln (30% formamide, 2x SSC, 0.06% triton)

2XSSC wash soln (2x SSC, 0.06% triton)

70%, 50%, 30% EtOH

#### Day1

Dissection:

- Standard dissection protocol, flies are usually 3-5 days old
- Dissected brains are fixed in 2% PFA at 25 ° C for 55 min
- Wash with 0.5% PBT, 3x, 10 min each

Dehydration:

- Dehydrate tissue with graded EtOH series: 30%, 50%, 70%, 100%, 100%, 100%, 10 min each
- Store the dehydrated tissue at 4 ° C with 100% EtOH overnight

#### Day2

Permeation:

- Rehydrate tissue with graded EtOH series: 70%, 50%, 30%, 10 min each
- Incubate tissue at 4 ° C 5% CH_3_COOH for 5 min (do this in the cold room if possible)
- Wash tissue with 4 ° C IxPBS, 3x, 5 min each
- Fix tissue with 2% PFA at room temperature for 55 min
- Wash tissue with 0.5% PBT, 3x, 10 min each

Autofluorescence quenching:

- Prepare 1% NaBH_4_ soln in 4 ° C 1xPBS (prepare fresh)
- Incubate tissue in 4 ° C with 1% NaBH4 for 30 min, change solution every 10 min (do this in the cold room if possible)
- Wash with 4 ° C IxPBS for 3x, 5 min each

Pre-Hybridization:

- Prepare fresh pre-hybr soln with Hi-Di™ formamide (Thermo Fisher Scientific)
- Incubate tissue with pre-hybr soln at 50 ° C for 2 hours

Hybridization:

- Take ~48uL hybr soln for 1 reaction. 3-5 brains per experiment.
- Add 1-2 uL of 50-100ng/uL of FISH probes to the hybr soln (total reaction value is 50uL)
- Incubate at 50 ° C for 10 hr, then at 37 ° C for 10 hr.

#### Day3

Washing:

- Warm up the pre-hybr soln to 37 ° C
- Add 200uL 37 ° C pre-hybr soln to the tissue with hybridization buffer, incubate for 10 min
- Wash with pre-hybr soln, 1x, 37 ° C, 10 min
- Wash with pre-hybr soln, 1x, at room temperature, 10 min
- Wash with 30% formamide, 2x, at room temperature, 30 min each
- Wash with 2xSSC washing soln, 3x, at room temperature, 10 min each
- Wash with 1xPBS, 1x, at room temperature, 10 min
- Fix tissue at 2% PFA at room temperature for 55 min
- Wash with 0.5% PBT, 3x, at room temperature, 10 min each

Clearing and mounting:

- Mount tissue on glass coated with poly-L-lysine
- Dehydrate with graded EtOH series: 30%, 50%, 75%, 100%, 100%, 100%, 10 min each
- Clear in xylene, 3X, 5 min each
- For confocal, mount in DPX (Electron microscopy sciences)

For BB-SIM, hold in xylene until imaging.

## Supplementary Note 3: Image Analysis

### Representation of BB-SIM data

Since the imaging plane of light sheet datasets collected by the BB-SIM microscope is typically tilted at a ~45° angle from the standard frontal/sagittal views, in some of the projections in the manuscript, the original BB-SIM stacks have been rotated and resliced to match standard sectioning conventions.

### Calculation of Cell intensity from Confocal Images

To quantify the fluorescence intensity of cells, we initially segmented individual cells using the Fiji^5^ (NIH) plugin TrakEM2^6^. We created subvolumes by manually tracing the cell bodies using the *Pdf* mRNA FISH signal as a reference. These same volumes were used to calculate integrated intensities by summing voxel intensities within each cell and subtracting the background level measured in adjacent voxels.

### Single Molecule Detection from BB-SIM stacks

For spot counting and intensity quantification purposes, individual fluorescent spots were automatically detected from BB-SIM z-stacks using a custom Matlab (Mathworks) algorithm^7^: the algorithm outputs the (*x*,*y*,*z*) position and intensity of each spot calculated by iterating a Gaussian mask algorithm^8^ (extended to 3 dimensions) on the dataset, after applying a local background correction. Since mRNAs are only a few tens of nm in size^9^, they are expected to appear as diffraction-limited particles. The dimensions of the 3D Gaussian mask were therefore chosen to match that of the Point Spread Function (PSF); the floating parameters were the (*x*,*y*,*z*) position and intensity of the spot. The background correction consisted in subtracting an affine fit of the intensities of the voxels adjacent to each spot. In order to calculate the number of spots per cell, we used 3D masks generated with TrakEM2^6^ around individual cell bodies (see above) and counted the spots which overlapped with each mask. Reconstructed images were generated using custom Matlab code that draws spots centered on each of the detected spots positions. Each spot is represented as a 3D Gaussian distribution which dimensions match that of the microscope PSF. We then generated depth-encoded projections of the 3D stacks using the Temporal Color Code command in Fiji^5^.

When quantifying the dimensions of individual spots, instead of the Gaussian mask algorithm we used custom Matlab code that fits the background-corrected intensity around each spot candidate using a PSF with adjustable size: we selected a cube-shaped region centered around each spot candidate extending ~3 PSF standard deviations in each direction (9x9x11 voxels). We first subtracted the local background by fitting the intensity within a 1-voxel thick region surrounding the 9x9x11 cube to an affine function of the 3D position. We then subtracted this estimate of the background from the intensity values in the cube. The resulting background-corrected intensity was fitted to a Gaussian PSF model using a Non-linear least square fit (Trust-region reflective algorithm), letting the (*x_0_*,*y_0_*,*z_0_)* position, Intensity (*I_0_*),. as well as PSF width (*σ_xy_*) and height (*σ_z_*) vary:

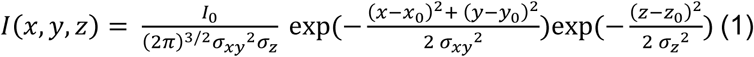

### Characterization of Single mRNA detection Sensitivity

In order to quantitatively assess the sensitivity and selectivity of our technique, we imaged z-stacks of PDF neurons (wild type flies, ZT14) labeled against *Tim* mRNA (Supplementary Figure S6a-e). We computed the fluorescence intensity distribution of detected spots in our z-stack using an intentionally low threshold to ensure that some of the brighter background pixels were included in the analysis. The resulting intensity distribution of detected particles is bimodal (Supplementary Fig. S6f), consisting of low-intensity background spots, and high intensity spots corresponding to the individual mRNAs. The bimodal intensity distribution is well fit by a sum of two lognormal distributions (Supplementary Fig. 6f, Equation 2); lognormal is the expected distribution resulting from the fluorescence intensity of individual molecules^10^.

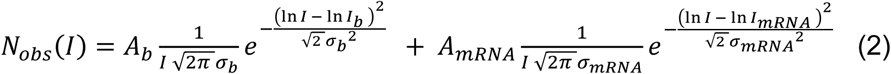

The two-component statistics provide a means to quantitatively predict the expected number of mRNAs vs. background spots detected as a function of a chosen intensity threshold (Equations 3-4):

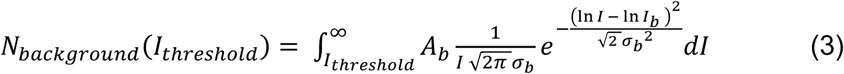

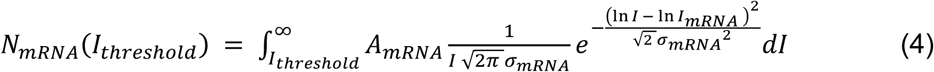

Using these estimates, we can quantitatively assess the similarity between the detected spots (*N_background_*(*I_threshold_*) + *N_mRNA_*(*N_threshold_*)) and the desired distribution *N_mRNA_*(0). We use the Jaccard Coefficient *J* as a metric^11^; the Jaccard Coefficient quantifies the similarity between two ensembles *A* and *B* as 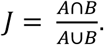 In our case:

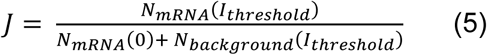

As expected, the Jaccard Coefficient reaches a maximum when the chosen intensity threshold is positioned at the shoulder between the two components of the bimodal distribution. The maximum value is excellent (*J* = 0.9; ~3% false positive detections; ~7% missed mRNAs; Supplementary Fig6f), indicating that the imaging and analysis provide an accurate absolute count of the number of individual mRNA molecules. As observed across the whole brain, the size or intensity of individual spots does not substantially vary with depth within the collected z-stack (Supplementary Fig. 6g).

### Quantification of *Pdf* mRNA localization along processes

In order to quantify the density of *Pdf* mRNAs along processes, we co-labeled brains with FISH probes against the *Pdf* gene and a HaloTag fluorescent ligand against a reporter myr::HaloTag fusion expressed in the PDF neurons. We acquired an initial low-resolution BB-SIM stack of the entire brain to provide the entire anatomical context (Fig. 3c), and then a second, highresolution stack, centered on the PDF neurons (Fig. 3d). We used the myr::HAloTag signal to trace processes that initiated from PDF neurons, using the Simple Neurite Tracer plugin in Fiji^5^. Only processes which origin could be unambiguously traced to a PDF neuron were considered. We then exported the path coordinates and used custom Matlab code to assign separately detected *Pdf* mRNA spots (see Single Molecule Detection from BB-SIM stacks section) to their relative positions along the neurite paths. Briefly, we selected FISH spots that lied within a maximum distance of 1 μm from the traced neurite path; we then computed for each selected FISH spot the distance from the cell body to the closest point along the neurite path (Fig.3e). We finally generated a FISH spot density histogram by computing the number of FISH spots detected at different distances from the cell body, normalized by the number of neurite path sections that mapped to these distances (Fig.3f). We obtained similar results when using a more stringent [neurite-FISH spot] distance of 500nm, or when using integrated FISH signal intensity instead of discrete spot counting as our metric.

